## Supplementary materials for "The effect of hybridization on transposable element accumulation in an undomesticated fungal species"

**Table S1. Description of the wild strains used in this study**

Table_S1.ods

CNs of full-length Ty1, Ty3p and Tsu4 (expressed as $\log_2$ NRD) are indicated for individual strains.


2

**Table S2. AIC values for linear models testing the association between population structure and environmental variation**

| Dataset | Family | AIC | |
|---|---|---|---|
| | | **gPCs+ePCs** | **gPCs** |
| All lineages | Ty1 | 562.75 | 569.19 |
| All lineages | Ty3p | 462.21 | 459.91 |
| All lineages | Tsu4 | 705.99 | 704.18 |
| *SpB* | Ty1 | 109.64 | 172.09 |
| *SpB* | Ty3p | 103.16 | 162.77 |
| *SpB* | Tsu4 | 294.08 | 306.74 |
| *SpC* | Ty1 | -30.77 | -2.88 |
| *SpC* | Ty3p | 33.13 | 33.85 |
| *SpC* | Tsu4 | 31.06 | 66.3 |


3

**Table S3. Nucleotide diversity at four-fold degenerate sites in protein-coding genes ($\pi_S$).**

| Lineage | $\pi_S$ |
|---|---|
| *SpA* | 0.0000334 |
| *SpB* | 0.0025064 |
| *SpC* | 0.0007916 |
| *SpC** | 0.0004162 |
| $SpD_1$ | 0.0000012 |
| $SpD_2$ | 0.0008641 |



4

**Table S4. Primers used for *ADE2* deletion in the parental *SpC* strains generated in this study.** Primers were described in Charron, Marsit et al., 2019.

| Name | Sequence |
|---|---|
| CLOP97-F5 | acaattaaggaatcaagaaaccgtgataaaaaattcaagtCAGCTGAAGCTTCGTACGC |
| CLOP97-F6 | gtaattgttcgctggccaagtatattaatacatttatataGCATAGGCCACTAGTGGATC |



5

**Table S5. Parental strains used in the MA experiment made in this study.**

| Background | Ploidy | Genotype | Reference |
|---|---|---|---|
| LL2011_001 | Haploid | *MATa hoΔ::kanMX4* | Charron et al., 2014 |
| LL2011_001 | Haploid | *MATa hoΔ::kanMX4 ade2Δ::hphNT1* | This study |
| LL2011_012 | Haploid | *MATa hoΔ::kanMX4* | Leducq et al., 2016 |
| LL2011_012 | Haploid | *MATa hoΔ::kanMX4 ade2Δ::hphNT1* | This study |
| MSH-587-1 | Haploid | *MATa hoΔ::natMX4* | Charron et al., 2014 |
| MSH-587-1 | Haploid | *MATa hoΔ::natMX4 ade2Δ::hphNT1* | This study |

**Table S6. Description of the MA lines used in this study.**

Table_S6.ods

CNs of full-length Ty1, Ty3p and Tsu4 (expressed as $\log_2$ NRD) are indicated for individual strains.

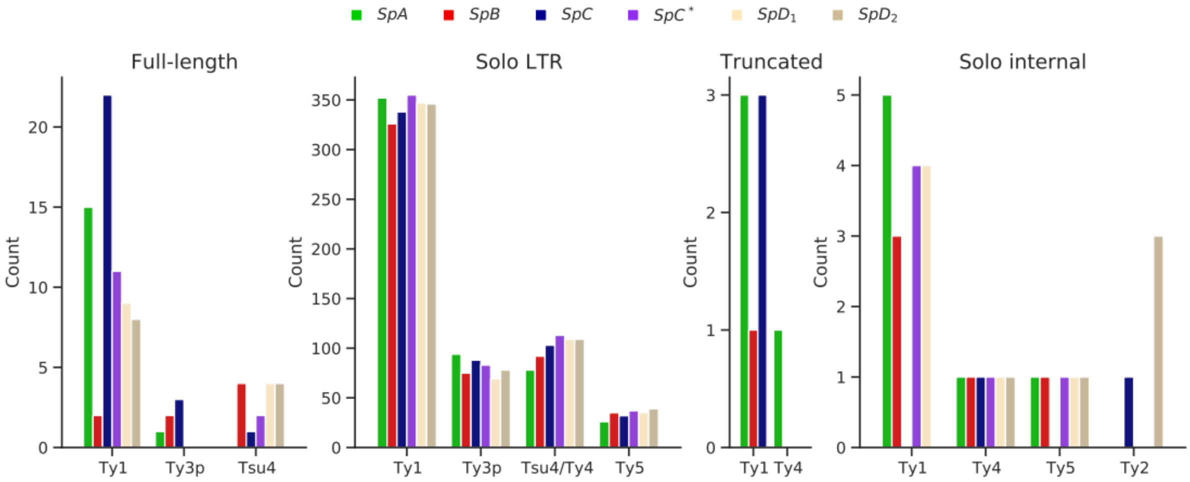


**Figure 2—figure supplement 1. Counts of LTR retrotransposon annotations in the six whole-genome assemblies.**


8

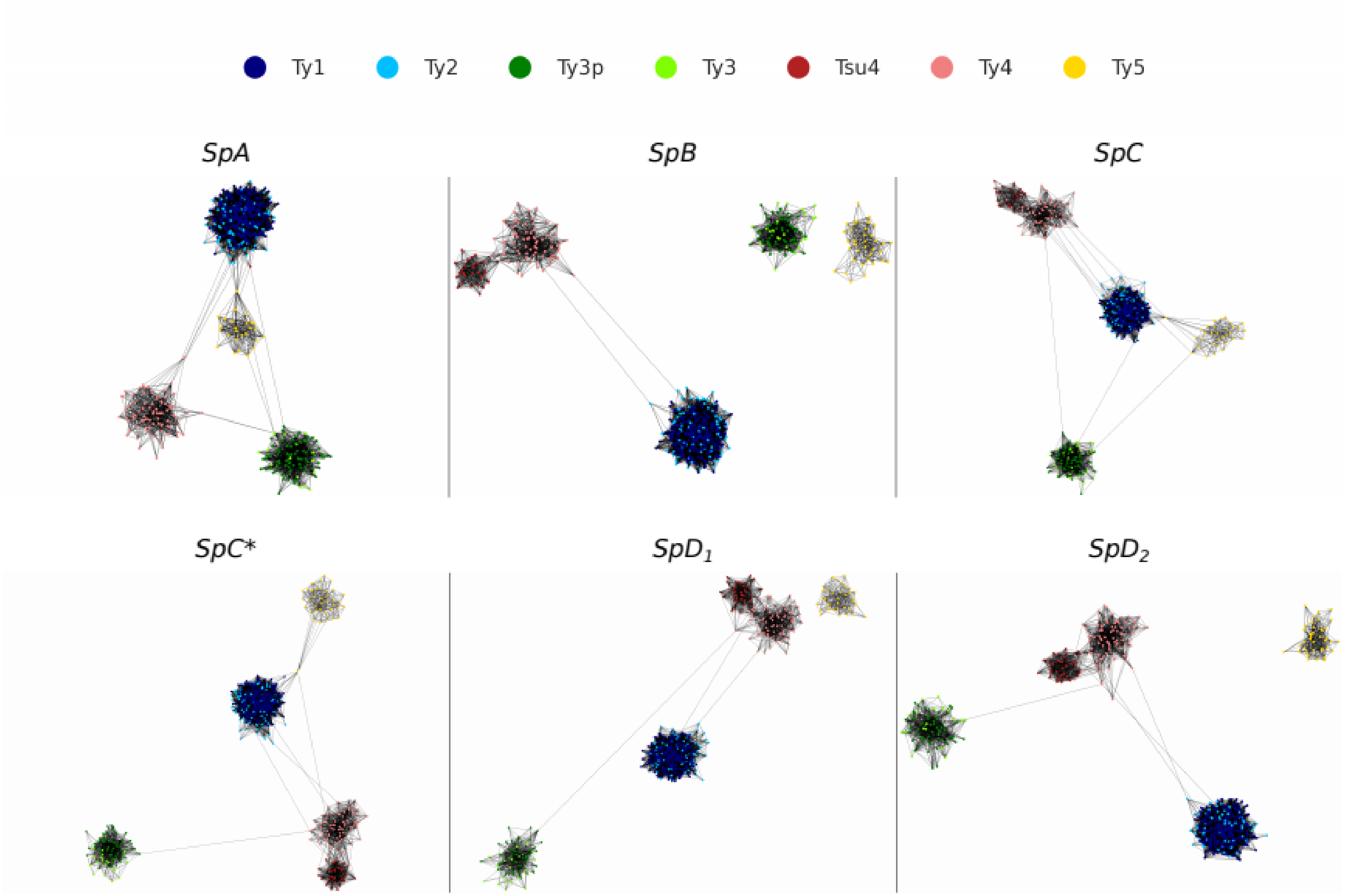


**Figure 2—figure supplement 2. LTR sequence similarity networks.** Networks were built using blastn bit scores as a similarity metric, keeping the top 10% interactions for each node. Networks were plotted using Cytoscape with the Prefuse Force Directed layout algorithm.



9

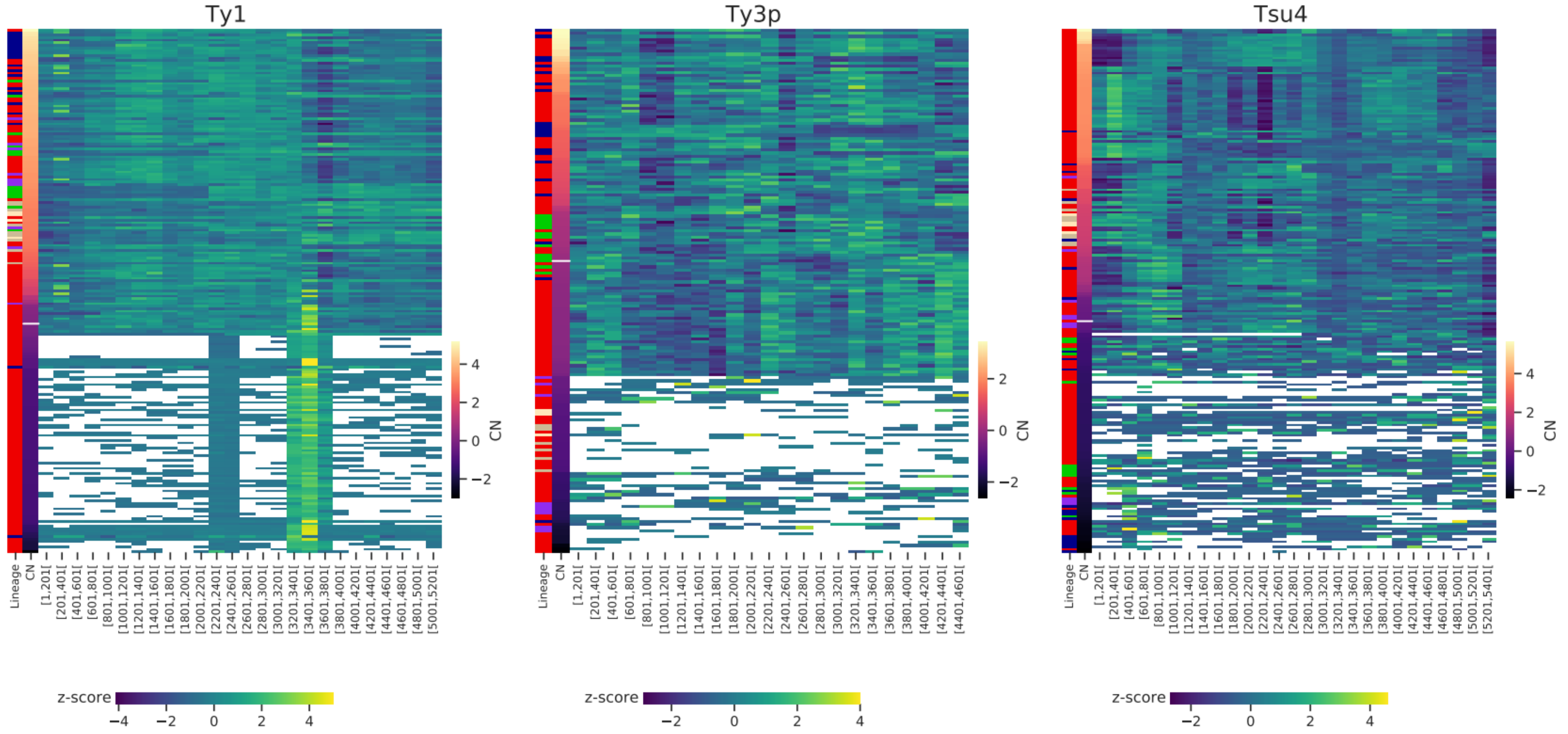


**Figure 2—figure supplement 3. Depth of coverage on reference internal sequences for Ty1, Ty3p and Tsu4.** Row z-scores are shown for 200 bp-wide non-overlapping windows. Phylogenetic lineage assignment and Ty CN ($log_2$ normalized read depth) are shown in the two first columns. White bars correspond to CN=0 (i.e. one copy).


10

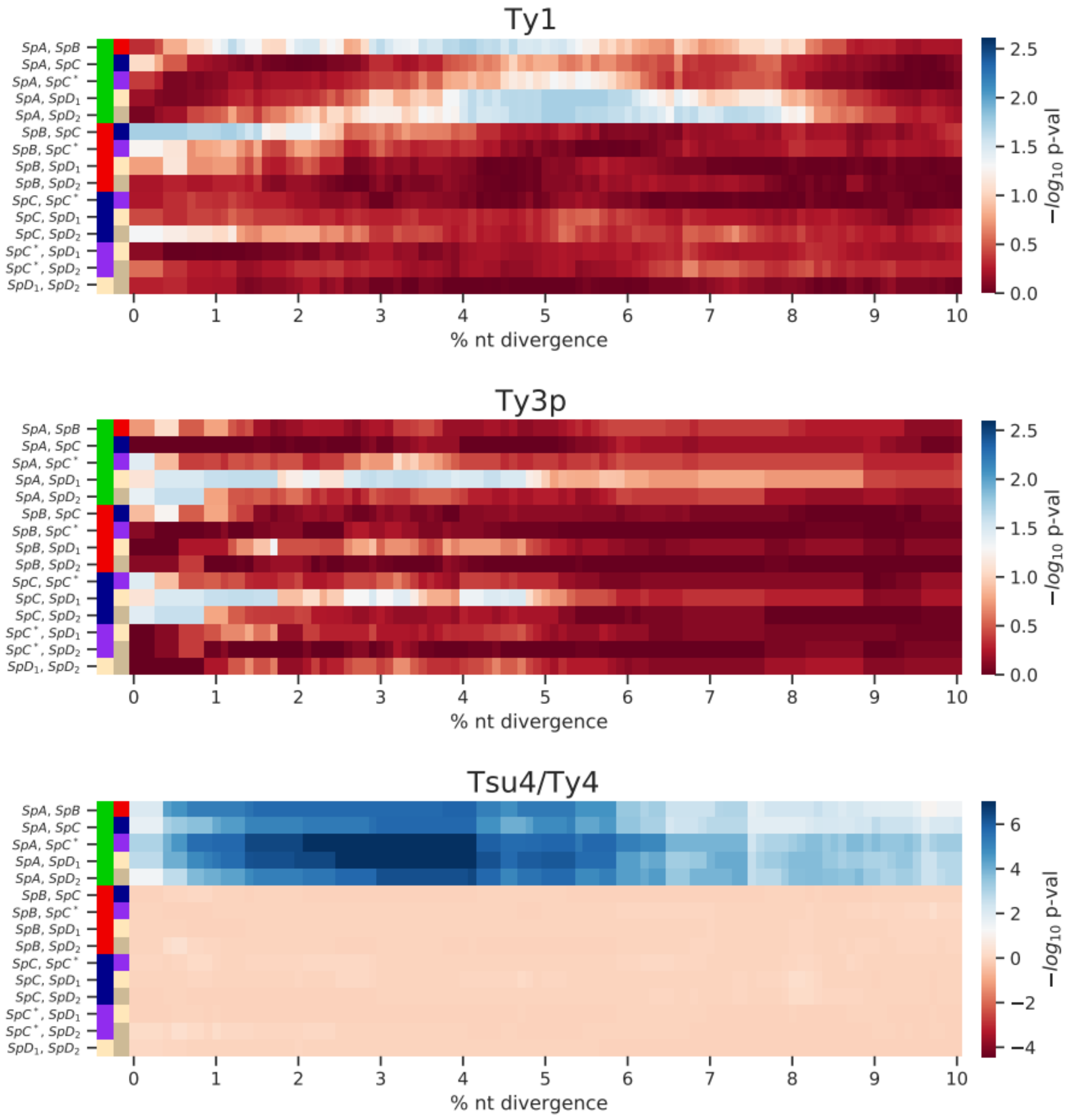


**Figure 3—figure supplement 1. Enrichment of low-divergence peaks in distributions of minimum nucleotide divergence between LTR sequences.** Heatmaps show FDR-corrected p-values for pairwise Fisher's exact tests between ratios of elements below and strictly above thresholds of nucleotide divergence from 0 to 10%. Color maps are centered at the significance threshold of 0.05.


11

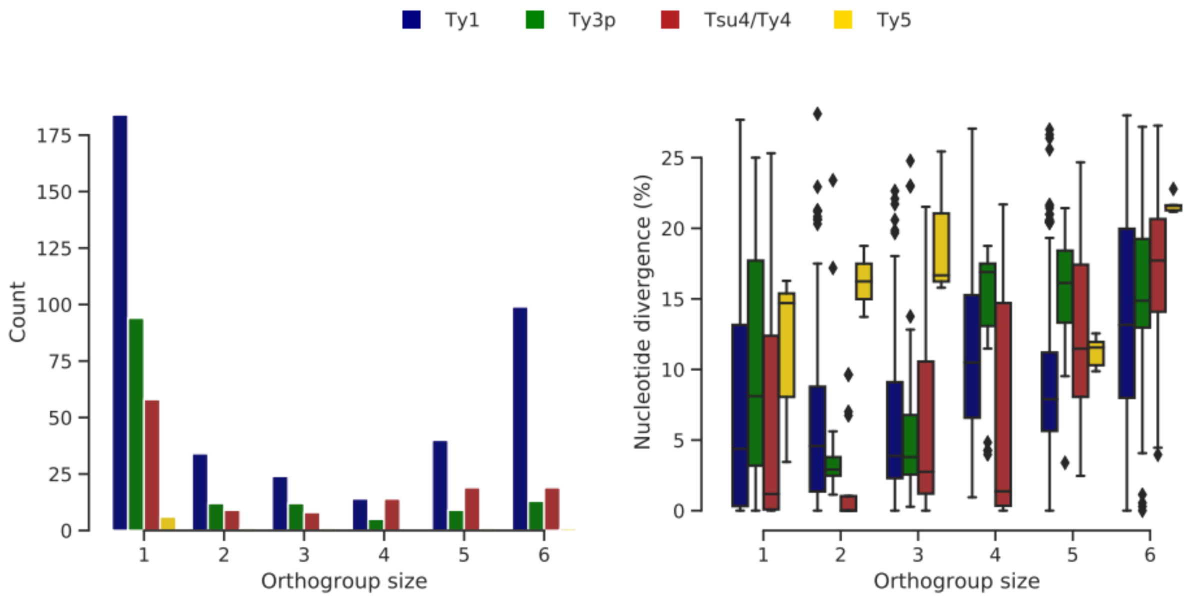


**Figure 3—figure supplement 2. Ty orthogroups defined from the six genome assemblies.** **a.** Distributions of Ty orthogroup sizes, from private (1) to conserved (6). **b.** Distributions of nucleotide divergence to the most closely related LTR sequence for each LTR within an orthogroup size category. Whiskers span 1.5 times the interquartile range.


12

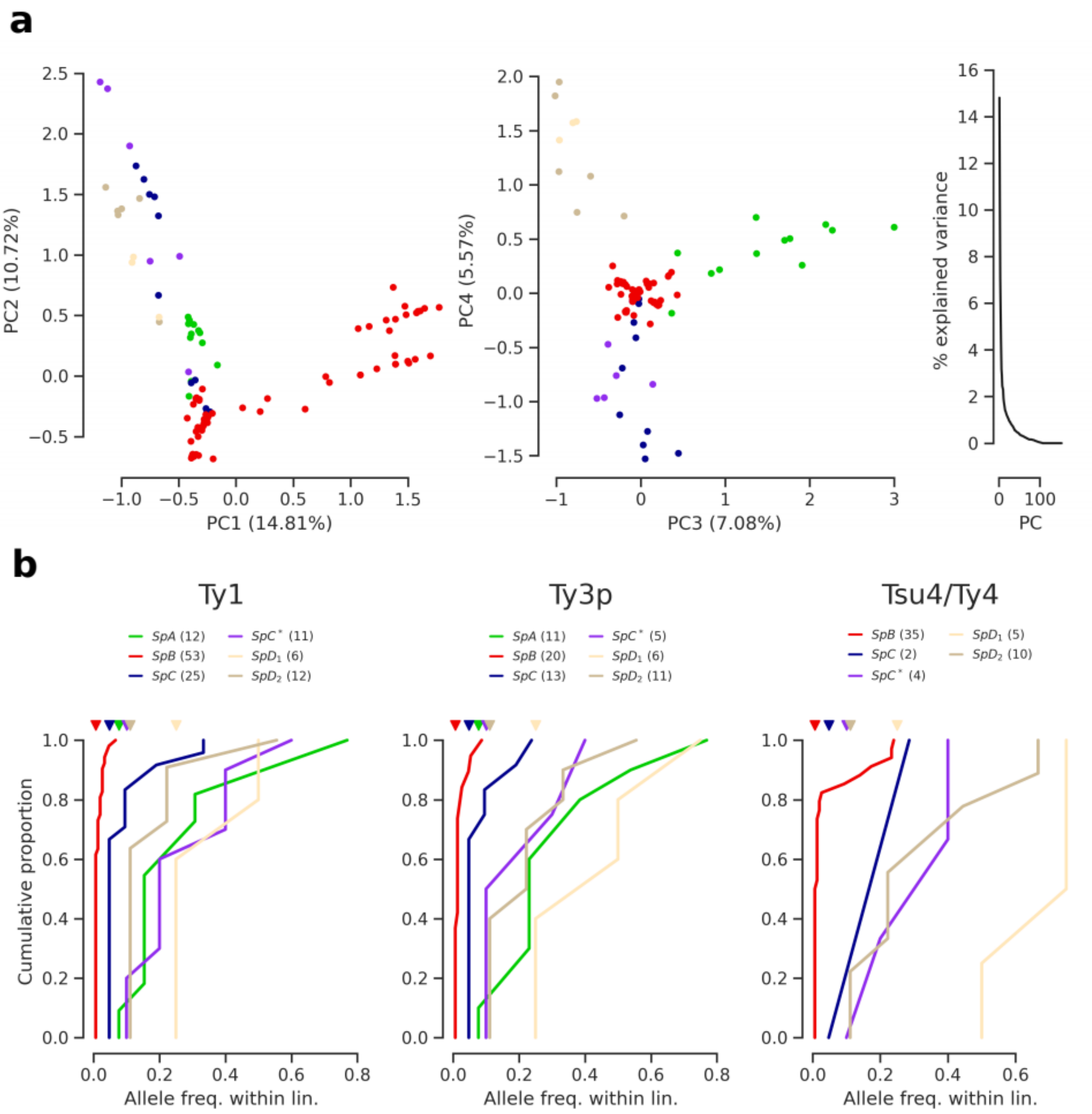


**Figure 3—figure supplement 3. Population structure based on Ty insertions called from discordant short read mappings. a.** PCA on Ty insertion calls showing the first four PCs and the distribution of explained variance. **b.** Cumulative distirbutions of Ty allele frequencies within lineages. Triangle marks on the top indicate allele frequencies corresponding to one strain. Numbers in parentheses correspond to counts of alleles.

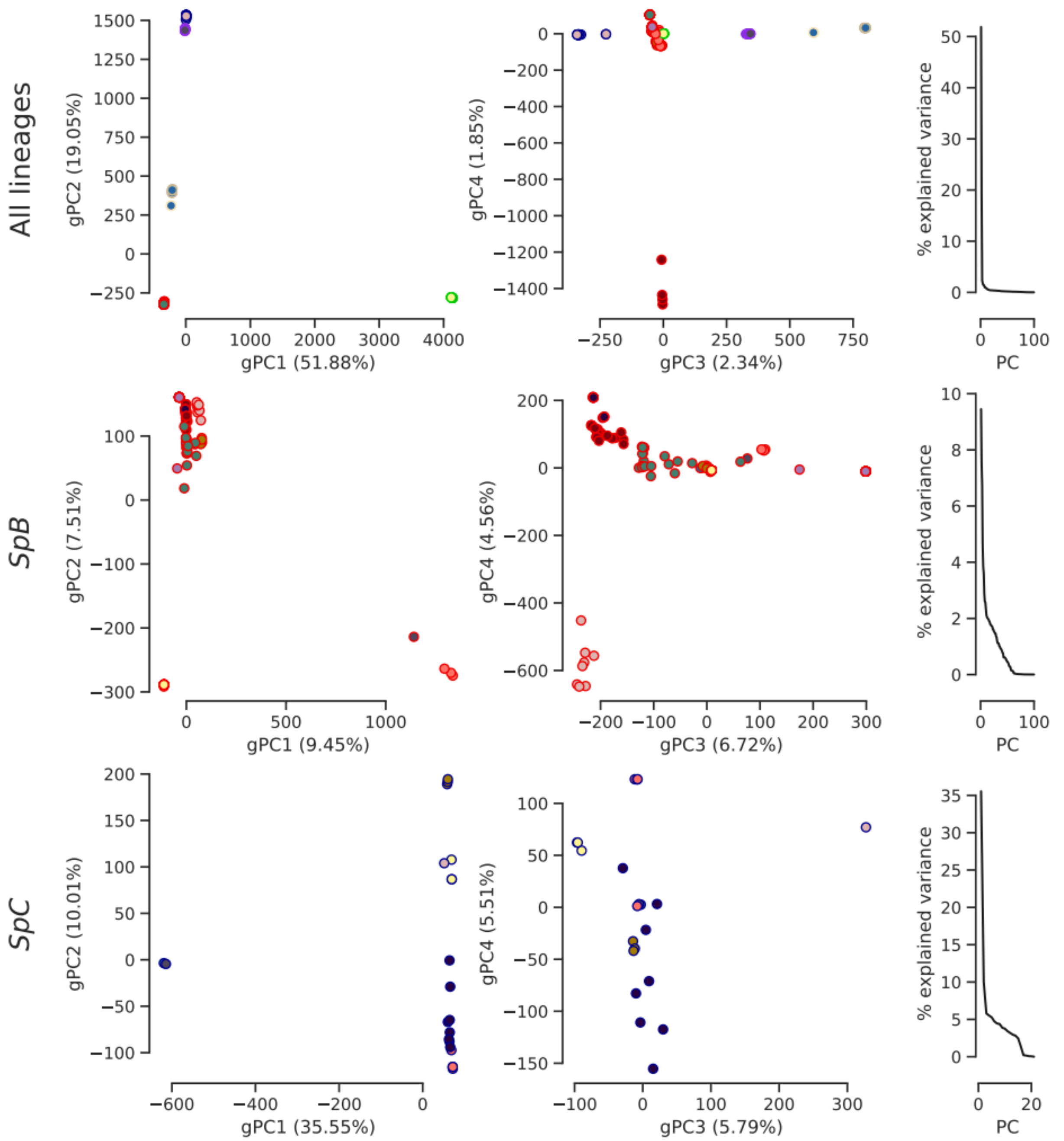


**Figure 4—figure supplement 1. Population structure based on genome-wide SNPs.** PCA on SNP calls showing the first four PCs and the distribution of explained variance. Dot face colors correspond to the clusters used in Figure 4b and Figure 4—figure supplement 4, and dot edge colors correspond to the phylogenetic lineage assignment.


14

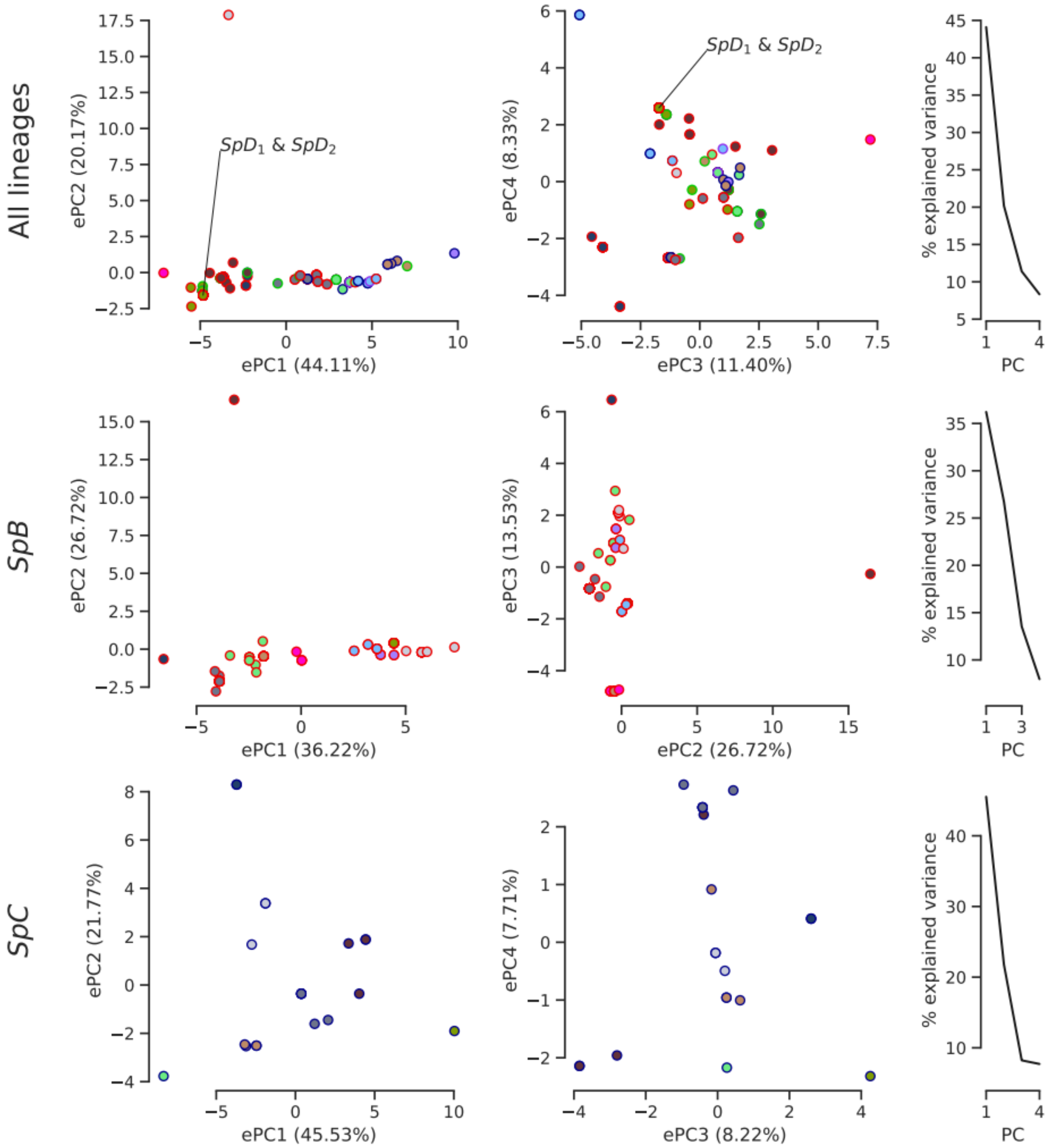


**Figure 4—figure supplement 2. Climatic variation among natural lineages.** PCA was performed on high-resolution climatic data (Abatzoglou *et al.* Sci Data 2018) at the sampling site of each strain. The first four or three PCs are plotted with the distribution of explained variance. Dot face colors correspond to the clusters used in Figure 4b and Figure 4—figure supplement 4, and dot edge colors correspond to the phylogenetic lineage assignment. The single sampling location of all *SpD* strains is labeled, as it overlaps with many *SpB* strains.


15

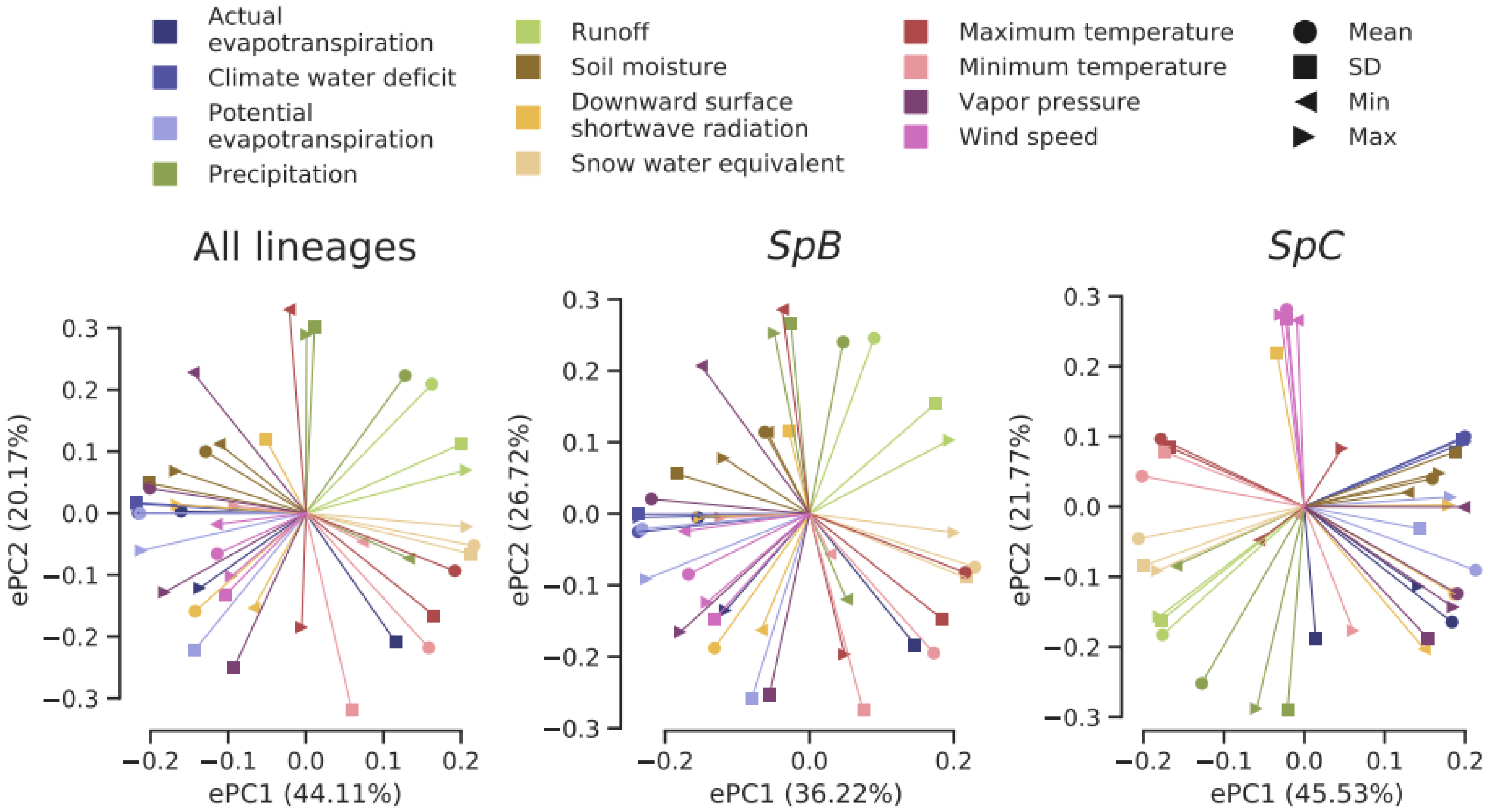


**Figure 4—figure supplement 3. Loadings of the PCAs on climatic variation data.** Eigenvectors associated with the two first PCs of each dataset are plotted.


16

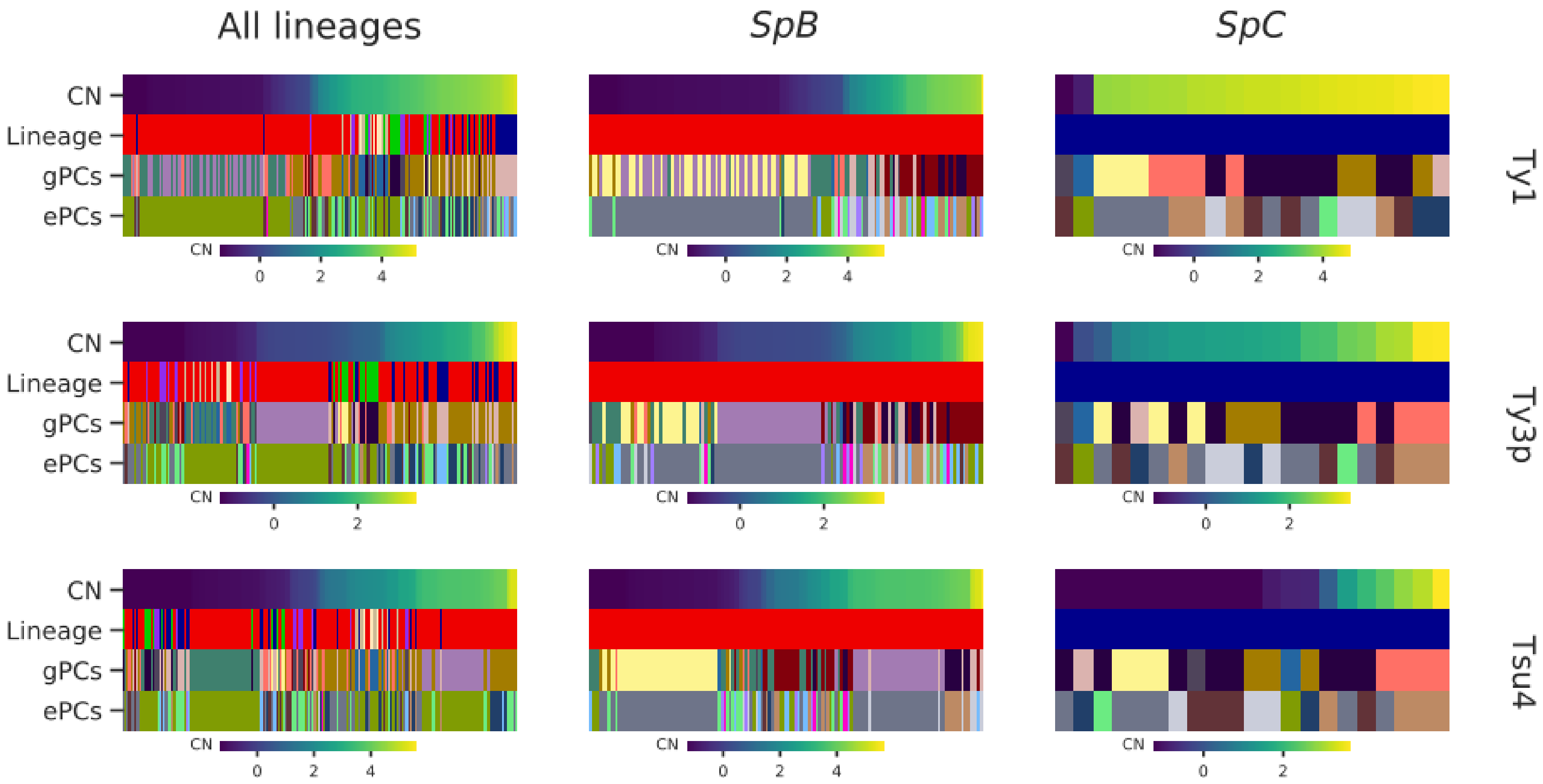


**Figure 4—figure supplement 4. Strains clustering performed on either gPC or ePC coordinates.** Columns within a heatmap correspond to individual strains. Ty CNs are represented by top rows, using the continuous color maps defined below each case. “Lineage” rows correspond to the established phylogenetic lineage assignment using the color code used throughout this paper. “gPCs” and “ePCs” correspond to clusters on gPCs and ePCs respectively, with discrete colors indicating the membership to different clusters. Cluster color mappings are consistent within datasets (All lineages, *SpB* and *SpC*).

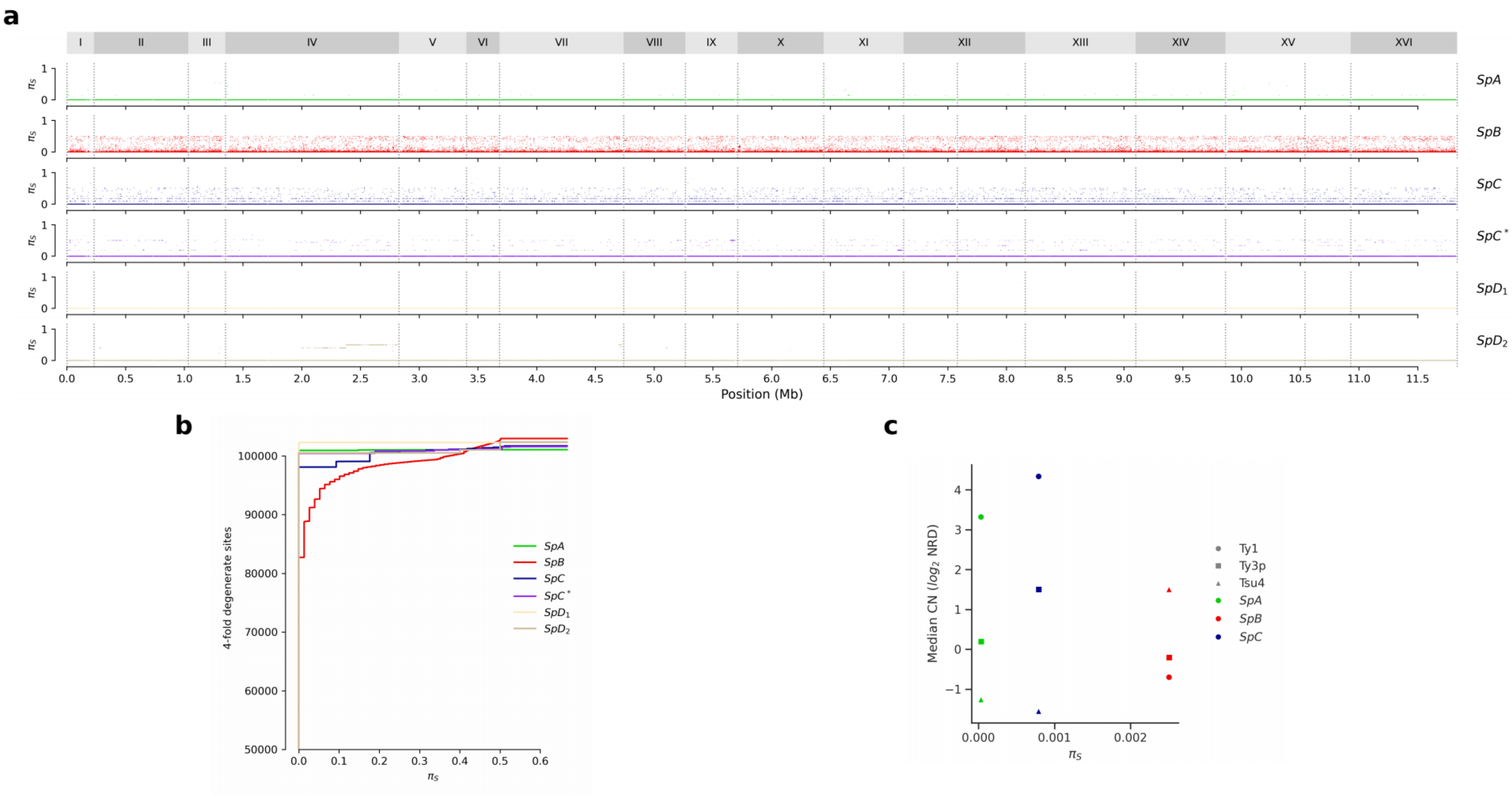


**Figure 5—figure supplement 1. Synonymous diversity at four-fold degenerate codon positions ($\pi_S$). a.** Genome-wide $\pi_S$. **b.** Cumulative distributions of $\pi_S$. **c.** Median Ty CN per lineage as a function of $\pi_S$.


18

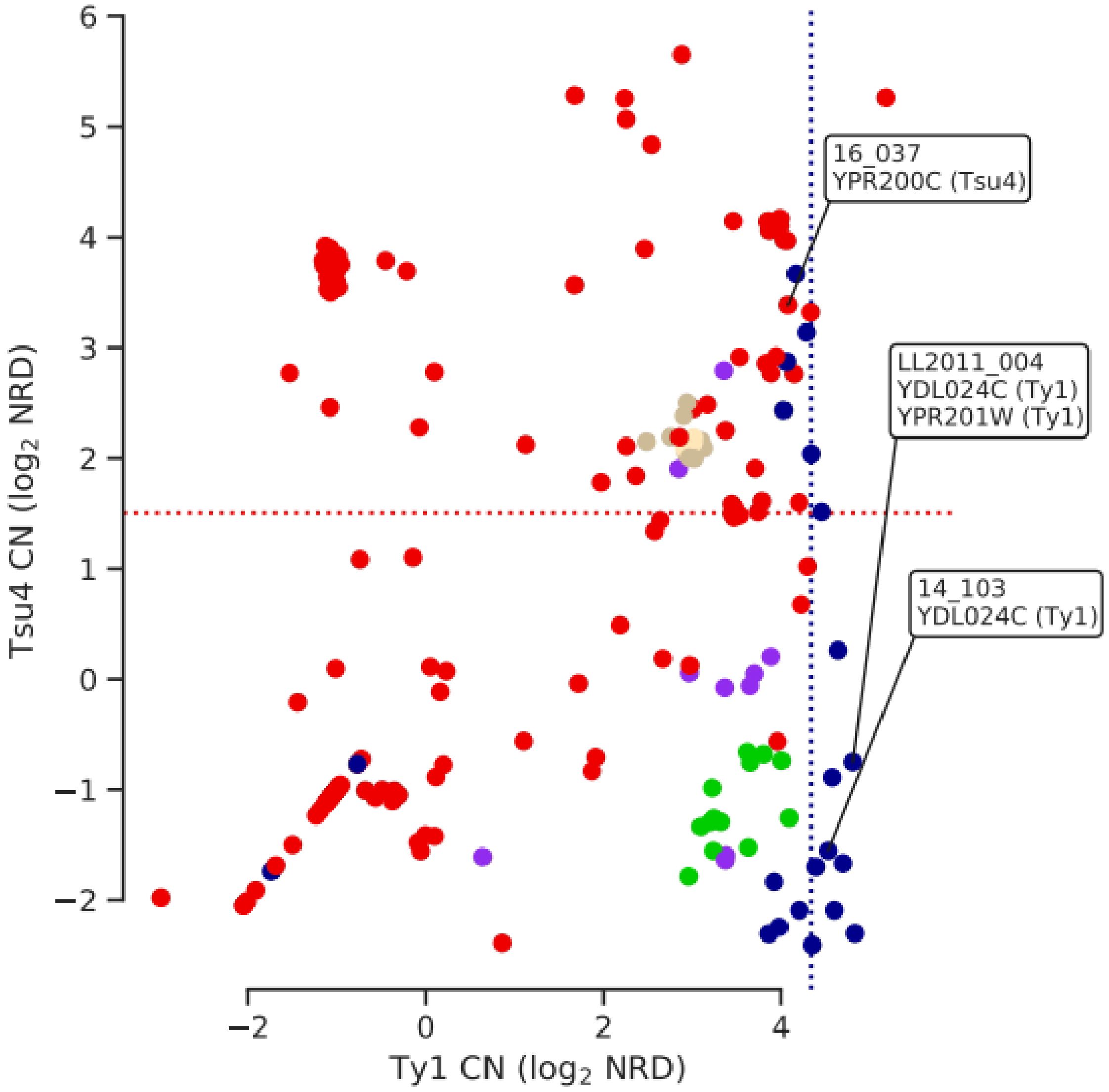


**Figure 5—figure supplement 2. Ty CNs of strains harboring ORF-disrupting insertions in relation with lineage-wide distributions of CN.** Strains with ORF-disrupting insertions are labelled with the corresponding gene and Ty family. Median Ty1 CN for *SpC* and median Tsu4 CN for *SpB* are shown as dotted lines.

 19

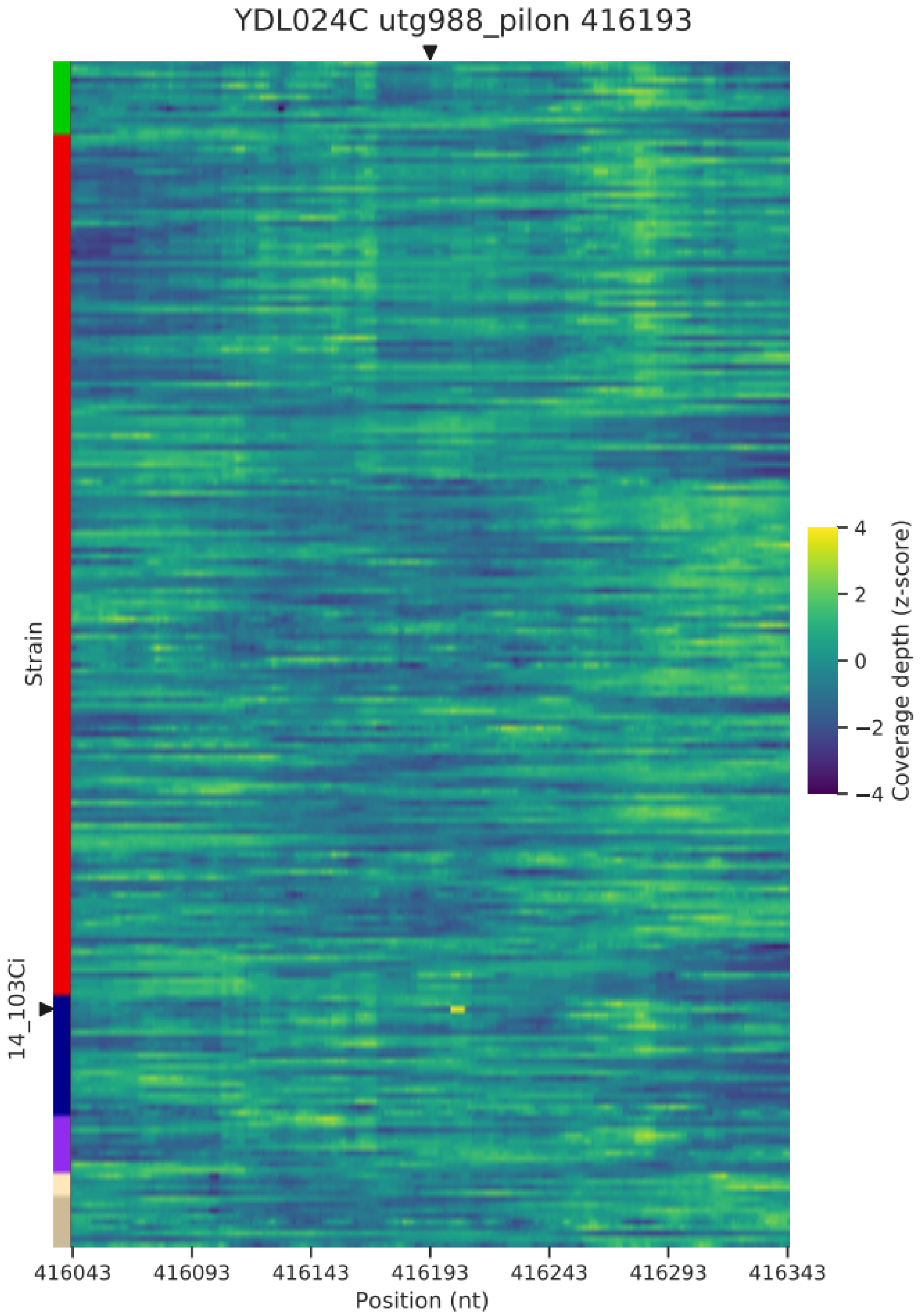


**Figure 5—figure supplement 3. Normalized depth of coverage at locus YDL024C-416.2 predicted to harbor Ty insertions.** Rows correspond to the 207 wild strains. Strains with predicted insertions are labelled on the y axes. Positions on the x axes correspond to 300 bp windows centered at the predicted locus.


20

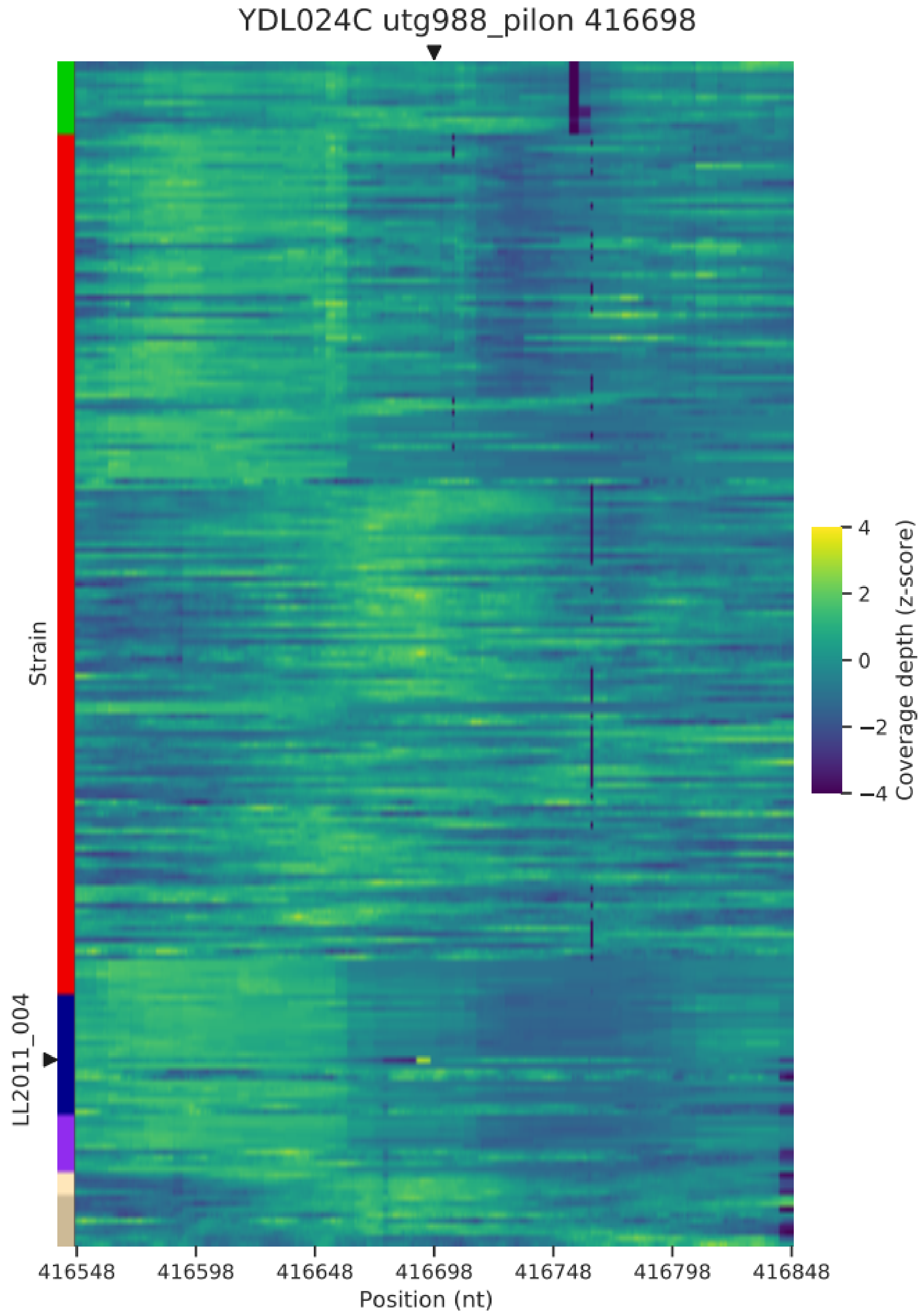


**Figure 5—figure supplement 4. Normalized depth of coverage at locus YDL024C-416.7 predicted to harbor Ty insertions.** Rows correspond to the 207 wild strains. Strains with predicted insertions are labelled on the y axes. Positions on the x axes correspond to 300 bp windows centered at the predicted locus.


21

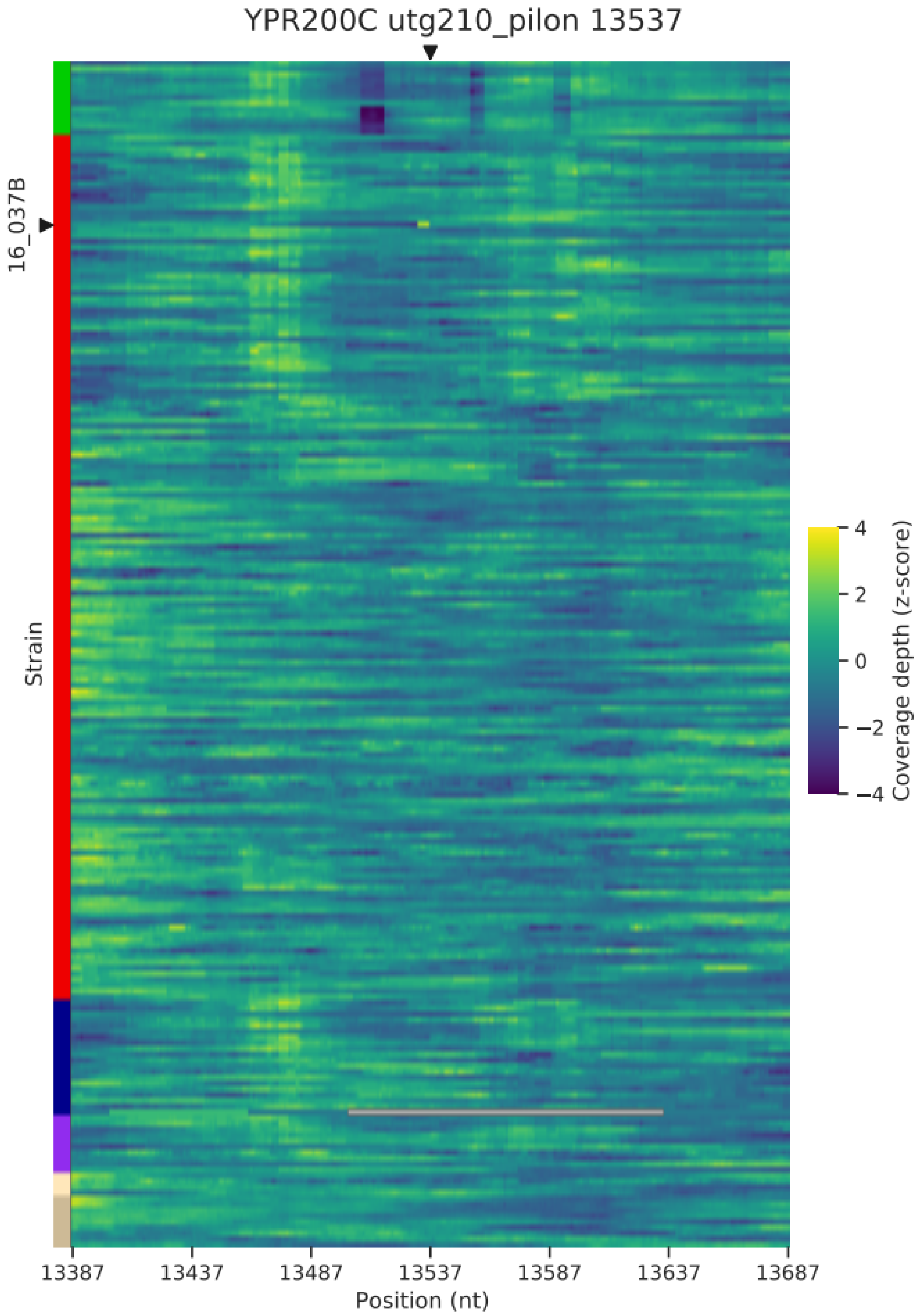


**Figure 5—figure supplement 5. Normalized depth of coverage at locus YPR200C predicted to harbor Ty insertions.** Rows correspond to the 205 wild strains. Strains with predicted insertions are labelled on the y axes. Positions on the x axes correspond to 300 bp windows centered at the predicted locus.

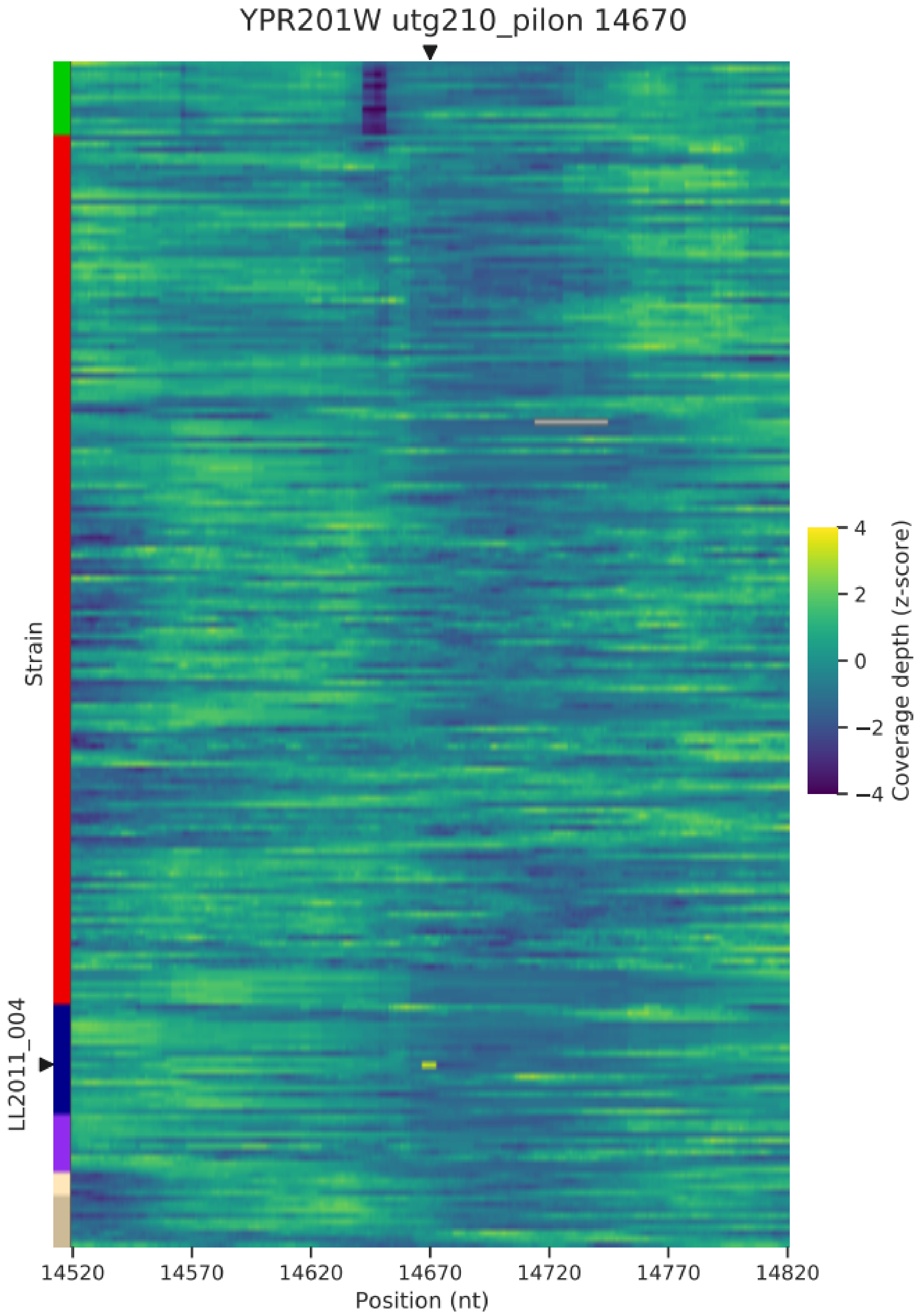


**Figure 5—figure supplement 6. Normalized depth of coverage at locus YPR201W predicted to harbor Ty insertions.** Rows correspond to the 206 wild strains. Strains with predicted insertions are labelled on the y axes. Positions on the x axes correspond to 300 bp windows centered at the predicted locus.

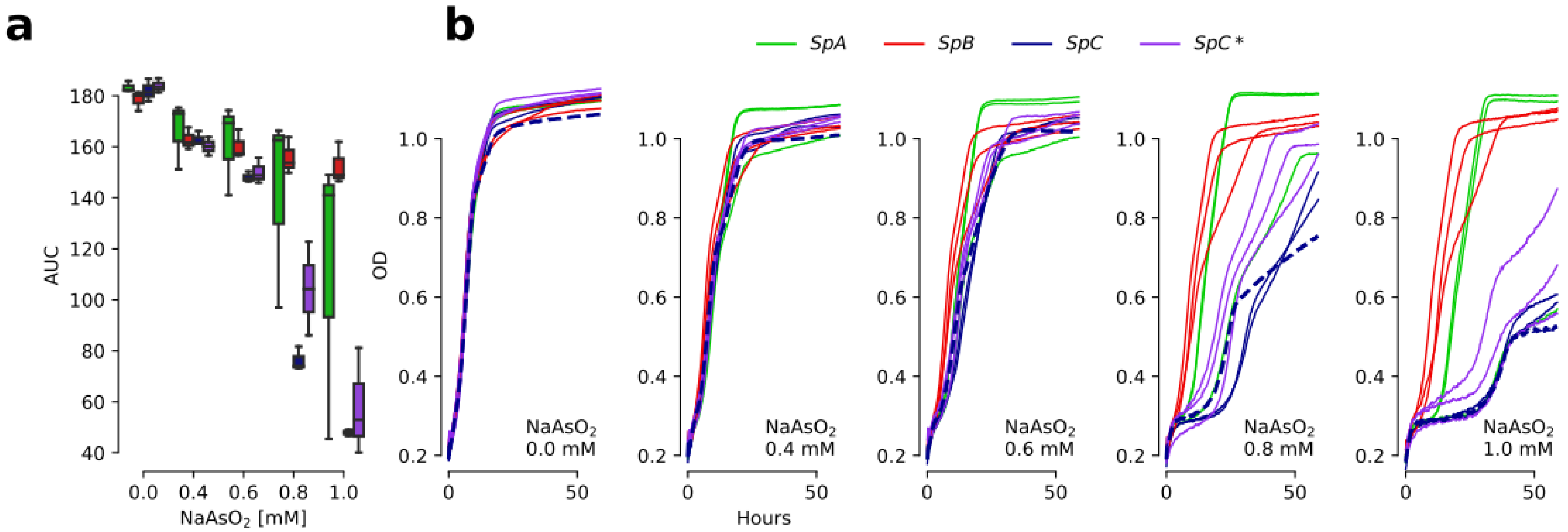


**Figure 5—figure supplement 7. Determination of the optimal $NaAsO_2$ concentration for growth measurements. a.** Growth curve area under the curve (AUC) at various concentrations of arsenite. Three representative strains per lineage were assayed. **b.** Growth curves showing optical density (OD) of cultures through time at 0, 0.4, 0.6, 0.8 and 1 mM $NaAsO_2$. The *SpC* strain LL2011_004 (which harbors a Ty1 insertion in *ARR3*) is highlighted with dotted curves. 0.8 mM $NaAsO_2$ was chosen as it best captures variation among *SpC* and *SpC** strains.

 24

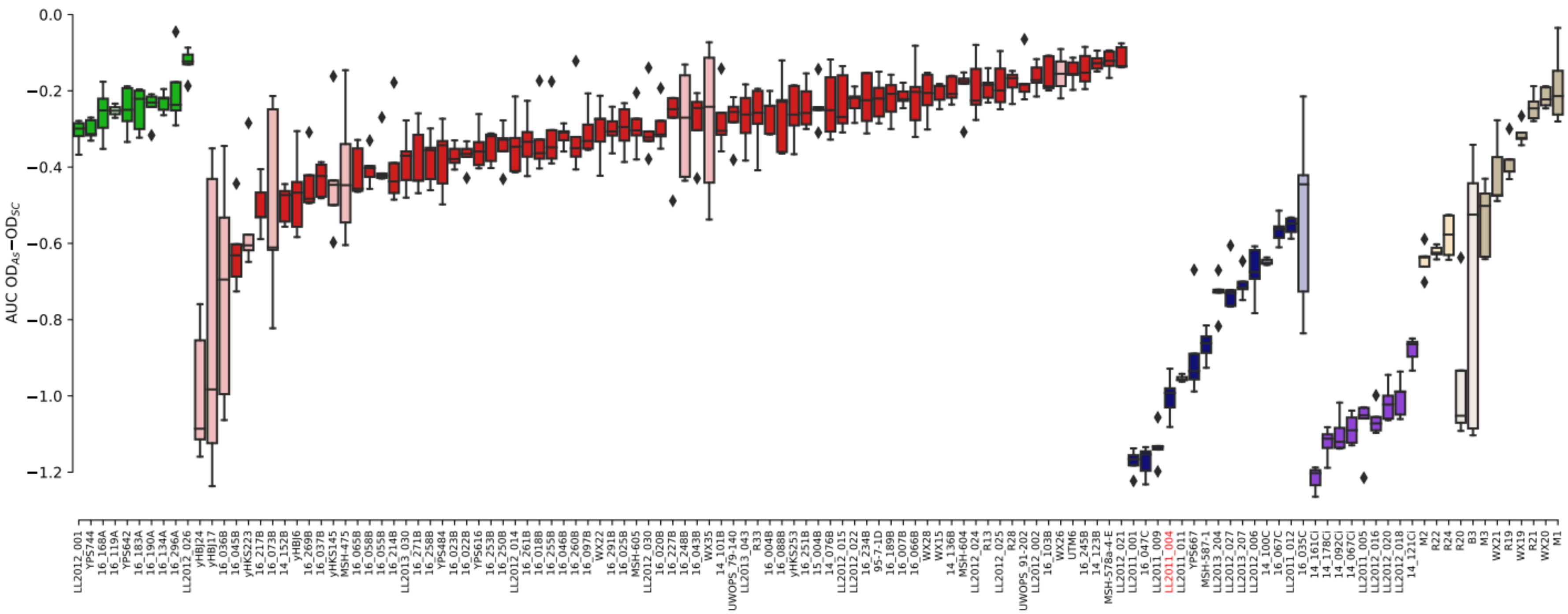


**Figure 5—figure supplement 8. Growth measurements of the collection of 123 strains in 0.8 mM $NaAsO_2$.** The difference between growth curves in 0.8 mM $NaAsO_2$ (As) and control medium (synthetic complete, SC) for individual replicates of each strain is plotted. Boxes in light color represent strains that were excluded from the linear models due to outlier variance or insufficient number of replicates (see Methods). Strain LL2011_004 is highlighted in red.


25

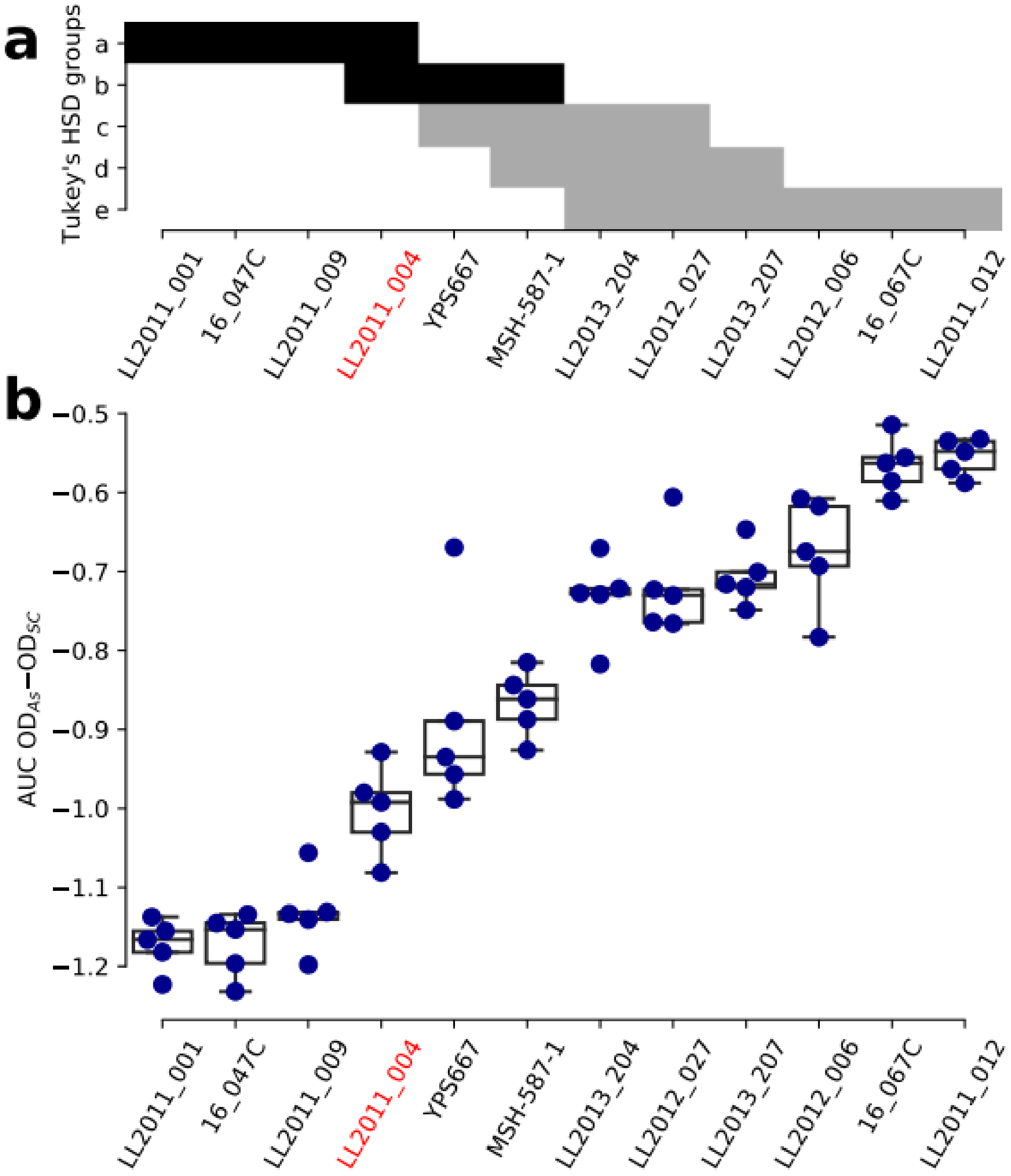


**Figure 5—figure supplement 9. Growth variation in $NaAsO_2$ within the *SpC* lineage. a.** Groups of strains that are not significantly different from a Tukey's HSD test on the AUC of the difference between growth in 0.8 mM $NaAsO_2$ (As) and control medium (synthetic complete, SC) for the *SpC* lineage. Groups comprising the strain LL2011_004 are shown in black. **b.** The AUC of the difference between growth curves in As and SC for the *SpC* lineage.

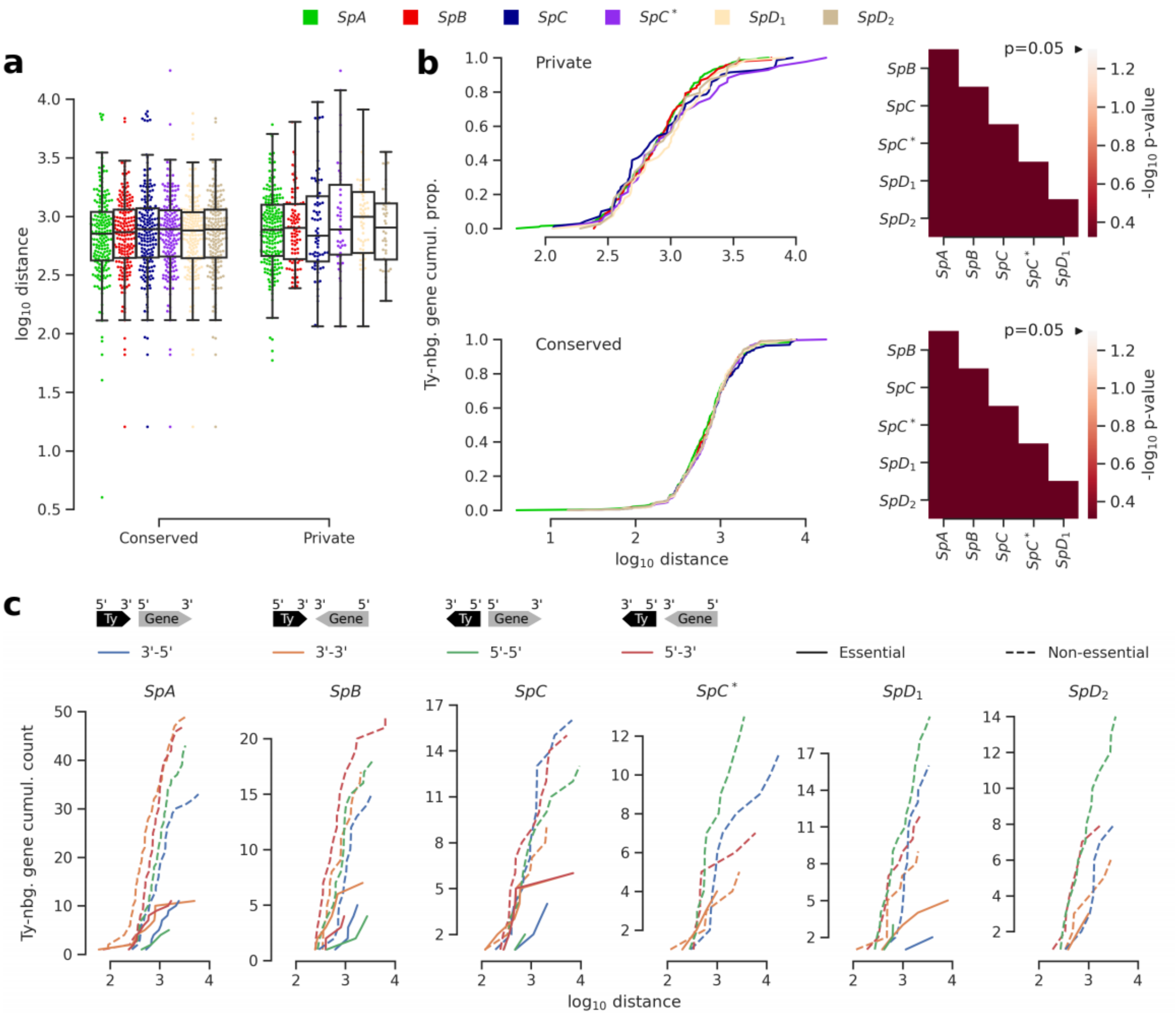


**Figure 5—figure supplement 10. Distance and orientation of Ty insertions relative to neighboring genes. a.** Distributions of distance to neighboring genes for Ty insertions conserved across the six genomes or private to one of them. **b.** Cumulative distributions of distance to neighboring genes for private (top) or conserved (bottom) insertions. Heatmaps show FDR-corrected p-values for pairwise Mann-Whitney U tests between distributions. Color maps are centered at the significance threshold of 0.05. **c.** Cumulative distributions of orientation of private Ty insertions relative to neighboring genes.

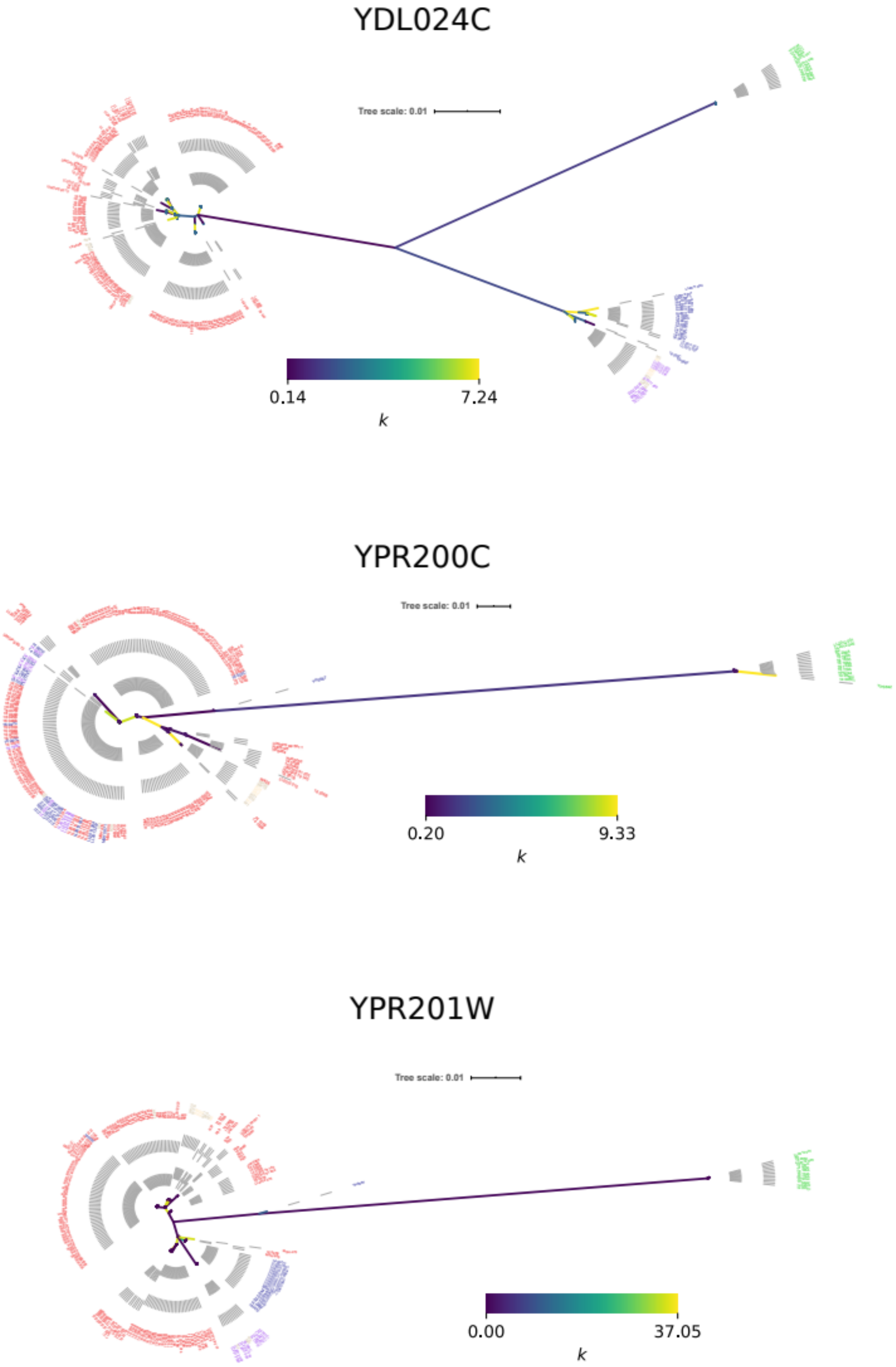


**Figure 5—figure supplement 11. Neighbor joining phylogenetic trees of genes with ORF-disrupting Ty insertions.** Branch color corresponds to the k parameter fitted by RELAX.

 28

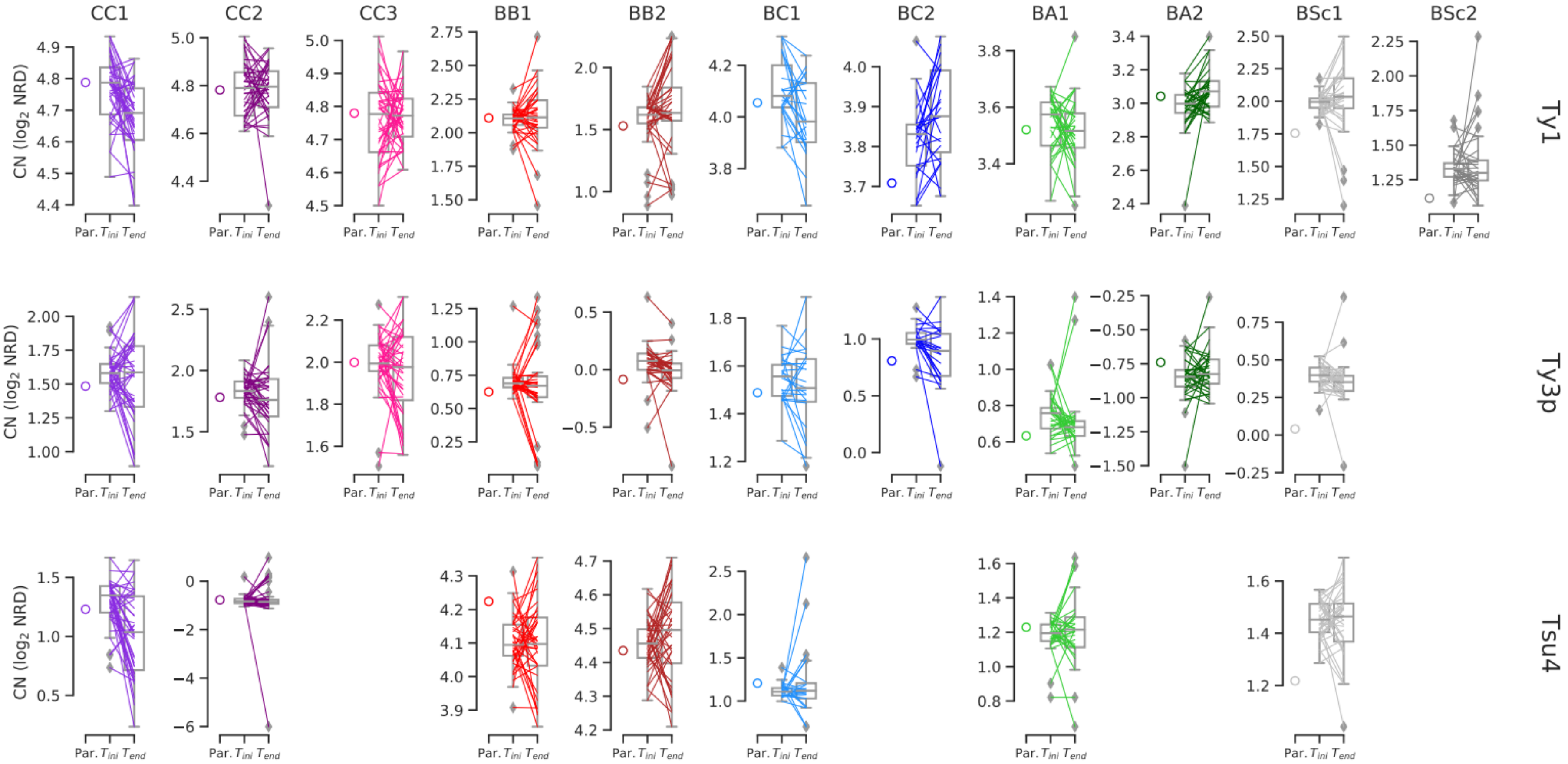


**Figure 6—figure supplement 1. Ty CNs in MA lines with predicted values for the combination of parental CNs.** CN values are the same as shown in Figure 6e, excepted that they are not normalized for median $T_{ini}$ CN. Whiskers span 1.5 times the interquartile range. Colored lines denote individual strains.

 29
